# IRBIT Directs Differentiation of Intestinal Stem Cell Progeny to Maintain Tissue Homeostasis

**DOI:** 10.1101/737262

**Authors:** Alexei Arnaoutov, Hangnoh Lee, Karen Plevock Haase, Vasilisa Aksenova, Michal Jarnik, Brian Oliver, Mihaela Serpe, Mary Dasso

## Abstract

The maintenance of the intestinal epithelium is ensured by the controlled proliferation of intestinal stem cells (ISCs) and differentiation of their progeny into various cell types, including enterocytes (ECs) that both mediate nutrient absorption and provide a barrier against pathogens. The signals that regulate transition of proliferative ISCs into differentiated ECs are not fully understood. IRBIT is an evolutionarily conserved protein that regulates ribonucleotide reductase (RNR), an enzyme critical for the generation of DNA precursors. Here, we show that IRBIT expression in ISC progeny within the *Drosophila* midgut epithelium cells is essential for their differentiation via suppression of RNR activity. Disruption of this IRBIT-RNR regulatory circuit causes a rapid, premature loss of intestinal tissue integrity as flies age. This age-related dysplasia can be reversed by suppression of RNR activity in ISC progeny. Collectively, our findings demonstrate an unexpected and novel role of the IRBIT-RNR pathway in gut homeostasis.

## Introduction

Like the mammalian intestinal epithelium, the *Drosophila* midgut epithelium is continually renewed by controlled ISC proliferation and differentiation of their progeny (Micchelli and Perrimon, 2006; Ohlstein and Spradling, 2006). ISC proliferation is finely tuned by diet, aging and the microbiota ecosystem (Choi et al., 2011; Koehler et al., 2017), using many of the same biochemical pathways that control intestinal epithelial renewal in mammals (Pasco et al., 2015). In addition to stem cell renewal, ISC division produces two types of postmitotic progeny: enteroendocrine cells (EECs) and enteroblasts (EBs). EBs ultimately mature into adult enterocytes (ECs) (Figure 1A). Mature ECs form the absorptive and protective surface of the epithelium (Micchelli and Perrimon, 2006; O’Brien et al., 2011; Ohlstein and Spradling, 2006; Zhai et al., 2017). Although ISC maintenance and proliferation has been extensively studied, the signals that mediate transition of ISC progeny into terminally differentiated absorptive ECs are not fully understood. The decision of ISC progeny to undergo differentiation is dictated by various intrinsic and extrinsic cues including nutrient availability and the presence of a physical damage in the intestine epithelium, and relies upon the level of interaction between ISC daughter cells. Daughters exhibiting low-level Notch signaling suppress Ttk69 transcriptional repressor and develop into EECs (Beehler-Evans and Micchelli, 2015; Wang et al., 2015; Zeng and Hou, 2015). Daughters with tight connections and strong Notch signaling commit to the EB lineage (O’Brien et al., 2011; Zhai et al., 2017). The process of terminal differentiation of the EB into the absorptive EC is not completely understood, but was shown to require the activity of several transcription factors, incuding Sox21a and GATAe (Zhai et al., 2017) (Figure 1A). The delay or block in terminal EC differentiation leads to accumulation of undifferentiated EBs, either causing dysplasia which can physically damage tissue integrity, or even neoplasia, with mosaic expression of various genes implicated in cancer progression (Hsu et al., 2014; Krausova and Korinek, 2014; Zhai et al., 2015).

**Figure 1.**
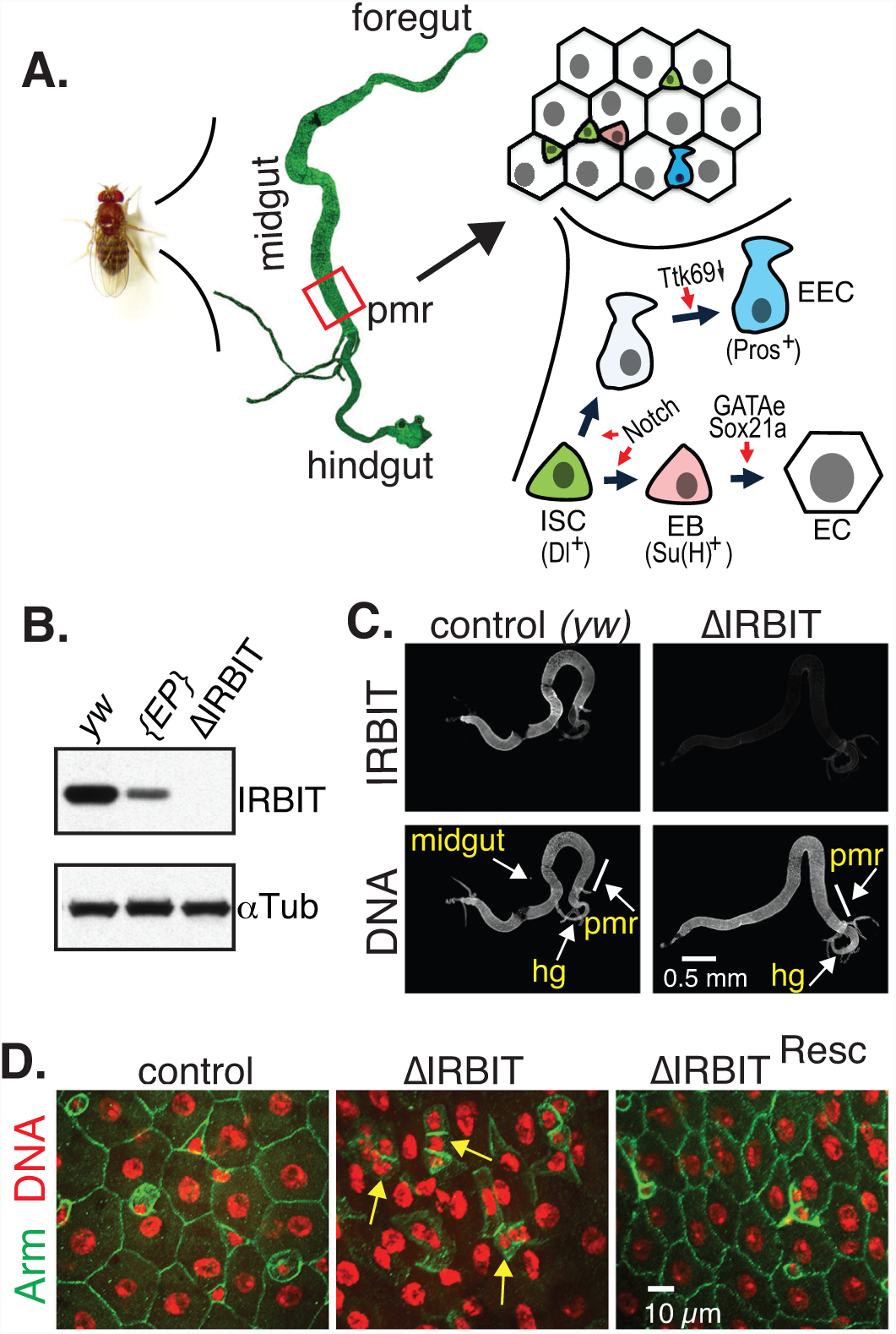
IRBIT is required for intestinal epithelial maintenance. (A) A scheme of digestive system in *Drosophila* and differentiation routes of intestinal stem cells (ISC) within posterior midgut region (pmr). EB (enteroblast), EC (enterocyte), EEC (enteroendocrine cell). (B) Total lysates of adult control (*yw*), P[EP]G4143 and ΔIRBIT flies were analyzed by Western blot for the presence of IRBIT. αTubulin was used as a loading control. (C) Guts of control and ΔRBIT flies stained with IRBIT antibodies and Hoechst 33342 (DNA). Posterior midgut region (pmr) is indicated. (D) Disruption of ΔIRBIT midguts architecture. Midguts of 12 d old control (*yw*) ΔIRBIT and ΔIRBIT^Resc^ flies stained for Armadillo (Arm, adherence junctions, green) and DNA (red). Arrows denote clusters of cells with small nuclei.

Ribonucleotide reductase (RNR) is a critical enzyme in the pathway for the de novo dNTP synthesis, as it makes dNDPs from corresponding ribonucleotide precursors via a remarkably complex mechanism (Ahluwalia and Schaaper, 2013; Fairman et al., 2011). RNR consists of two subunits: the R2 subunit provides the free radical that is necessary for R1 subunit-mediated reduction of ribonucleotides. In addition to the catalytic site, R1 subunit has two nucleotide binding sites that control the state of R1, and one of them, the A-site, monitors R1’s overall activity. RNR is active when ATP binds to the A-site, while dATP binding inhibits the enzyme (Ahluwalia and Schaaper, 2013). As the A-site has low affinity for ATP/dATP, the concentrations of dATP required to inhibit RNR usually exceed physiological dATP levels inside dividing cells. We have previously shown that an evolutionarily conserved protein IRBIT (IP_3_-receptor-binding protein released with inositol 1,4,5-trisphosphate) controls RNR activity by locking the R1 subunit in an R1*dATP inactive state, in the presence of physiologically relevant dATP concentrations (Arnaoutov and Dasso, 2014). While the dNTP pool in HeLa cells is sensitive to IRBIT levels, the organismal importance of IRBIT-dependent RNR regulation remained unknown although we speculated that it could control the cell cycle progression and exit (Arnaoutov and Dasso, 2014). High dNTP levels, produced by RNR, are critical for cells to transit through S phase. During *Drosophila* embryogenesis, the maternal pool of dNTP is only sufficient for the first 10 divisions, after which endogenous RNR activity becomes indispensable (Djabrayan et al., 2019; Song et al., 2017). On the other hand, overexpression of RNR appears to be detrimental for normal progression of embryogenesis (Song et al., 2017), suggesting that there must be mechanisms to curtail RNR activity during cellular specialization.

Because dNTP abundance is critical for S phase progression and because the suppression of the cell cycle could regulate differentiation (Djabrayan et al., 2019; Jiang and Kang, 2003; Ruijtenberg and van den Heuvel, 2016; Vastag et al., 2011), we decided to test whether manipulation of the RNR activity could affect cell’s choice between proliferation and differentiation.

Here, we tested the function of IRBIT in tissue homeostasis, particularly the proliferation and differentiation of ISCs and maintenance of the adult *Drosophila* midgut epithelium. We found that the IRBIT-RNR pathway is essential to ensure correct differentiation of ISC progeny. We show that conserved transcriptional factor GATAe stimulates IRBIT expression in postmitotic ISC progeny to inhibit RNR and promote differentiation. The intestines of flies lacking IRBIT demonstrate dysplasia, with profound accumulation of undifferentiated ISC progeny. Additionally, we show that the GATAe-IRBIT-RNR pathway may become dysfunctional as flies age, resulting in characteristic accumulation of undifferentiated ISC progeny. Such dysplasia can be successfully reversed by specifically inhibiting RNR in the ISC progeny. Collectively, these findings show that suppression of RNR activity by IRBIT is an indispensable mechanism in the ISC daughter cell to direct the latter towards differentiation and to maintain intestinal tissue homeostasis.

## Results

### IRBIT is required for intestinal epithelial maintenance

There are two *Drosophila* genes that encode proteins with significant sequence similarity to vertebrate *IRBIT*, *AhcyL1* (*CG9977, IRBIT*) and *AhcyL2* (*CG8956, IRBIT2*). Only IRBIT but not IRBIT2 bound RNR efficiently (Figure S1A), suggesting that IRBIT controls RNR in *Drosophila*. Notably, only IRBIT but not IRBIT2 mRNA was expressed in the midgut (Figure S1B). Thus, both protein-protein interactions and localized expression prompted us to focus on IRBIT control of RNR regulation in the midgut. We generated two null alleles of IRBIT and we termed flies bearing both as “∆IRBIT” (Figure S1C). We stained digestive tracts isolated from adult female flies with anti-IRBIT antibodies, confirming IRBIT protein expression in the midgut, as well as its absence in ∆IRBIT flies (Figures 1B and 1C). To verify the specificity of IRBIT loss-of-function phenotypes, we introduced a genomic rescue fragment to the defined docking site in the ∆IRBIT background and termed these flies as “∆IRBIT^Resc”^. ∆IRBIT flies expressed non-functional RNA and lacked IRBIT protein, while ∆IRBIT^Resc^ flies expressed IRBIT RNA at levels similar to controls (Figures S1D and S1E).

We focused on the function of IRBIT in the female posterior midgut region (pmr, R5 region (Dutta et al., 2015)) because this tissue has a well-characterized and relatively simple structure (Micchelli and Perrimon, 2006; Miguel-Aliaga et al., 2018; O’Brien et al., 2011; Ohlstein and Spradling, 2006; Zeng et al., 2010; Zeng and Hou, 2015; Zhai et al., 2017) (Figure 1A). By 12 d post-eclosion, the ∆IRBIT pmr epithelium showed degenerate tissue with hyperplastic-like polyps (Figures 1D, S2A and S2B), which consisted of cells with small nuclei instead of large differentiated ECs. The midguts of ∆IRBIT^Resc^ flies were normal (Figure 1D), confirming that the defects seen in ∆IRBIT midguts result from IRBIT loss. Although the lifespan of both ∆IRBIT and ∆IRBIT^Resc^ flies were similar (Figure S2C), EM ultrastructural analysis revealed that ∆IRBIT midguts have thinner peritrophic membrane (PM) – an extracellular matrix barrier against microbial infection (Figure S2D). We examined peritrophic membranes from the midguts of 8 d old axenic (free from microorganisms) flies by staining with lectin-HPA (Helix pomatia agglutinin). The PMs of ∆IRBIT’s midguts was clearly thinner than PMs of control or ∆IRBIT^Resc^ flies (Figure S2E), consistent with the EM ultrastructural analysis. Altogether, our results indicate that the loss of IRBIT in flies leads to formation of intestine that has weak PM and demonstrate tissue dysplasia.

### IRBIT mediates differentiation of the ISC progeny

∆IRBIT midgut dysplasia could result from increased ISC proliferation, failed transition of ISC progeny into EECs and EBs, and/or failed maturation of EBs into ECs. To test these possibilities, we determined the relative abundance of these cell types (Figures 2A and S3). Escargot (Esg) has a well-defined pattern in the midgut and is expressed in both ISCs and EBs, while the expression of Su(H)GBE is restricted to EBs. Delta (Dl) is a known marker of ISCs and Pros faithfully marks cells of EEC lineage (He et al., 2018). To enhance our arsenal of detection tools, we additionally used antibodies against several proteins that play key functions during the cell cycle and found that antibodies against Asterless (Asl, centriole component) uniformly stain EBs and ISCs, while antibodies against Polo (Polo kinase, Plk) specifically detect ISCs. Antibodies against R1 (RnrL), the large subunit of RNR, revealed that R1 is specifically expressed in both ISCs and EBs. (Figure S3). Midguts of ∆IRBIT flies had normal levels of ISCs, increased numbers of the ISC progeny – EBs and immature EECs – and reduced population of ECs. This pattern would be consistent with a block or delay in the differentiation of ISC progeny (Figures 2A, 2B, S4A and S4B). Clusters of small cells in 8d old ∆IRBIT pmr typically consisted of a single ISC and 2-4 attached undifferentiated progeny (Figure 2C). Moreover, these clusters invariably showed high levels of RNR by immunostaining (Figure 2D), suggesting that ISC progeny in ∆IRBIT midguts expressed high levels of RNR, failed to separate from mother ISCs and failed to differentiate. By contrast, we did not observe EB groups associated with ISCs in pmr of 8d old control flies, indicating that EBs detach from mother ISCs and differentiate rapidly, maintaining normal homeostasis (Figures 2B, 2C and 2D). Importantly, these R1^+^ clusters in ∆IRBIT midguts develop rapidly (as early as day 8 post eclosion) because we did not detect significant accumulation of R1^+^ cells in midguts of newborn ∆IRBIT flies (1d post eclosion) (Figure S4C). To visualize IRBIT expression during ISC-EB-EC transitions, we used a ^*ts*^IRBIT promoter (Figure 2E) to express nlsGFP (nuclear GFP) and examined the ISC niche. We found that nlsGFP was not detected in ISCs but accumulated in ISC progeny that were committed to differentiation, and remained highly expressed during the EB-EC transition (Figures 2E and S4D). This pattern suggested activation of IRBIT expression in the differentiating progeny.

**Figure 2.**
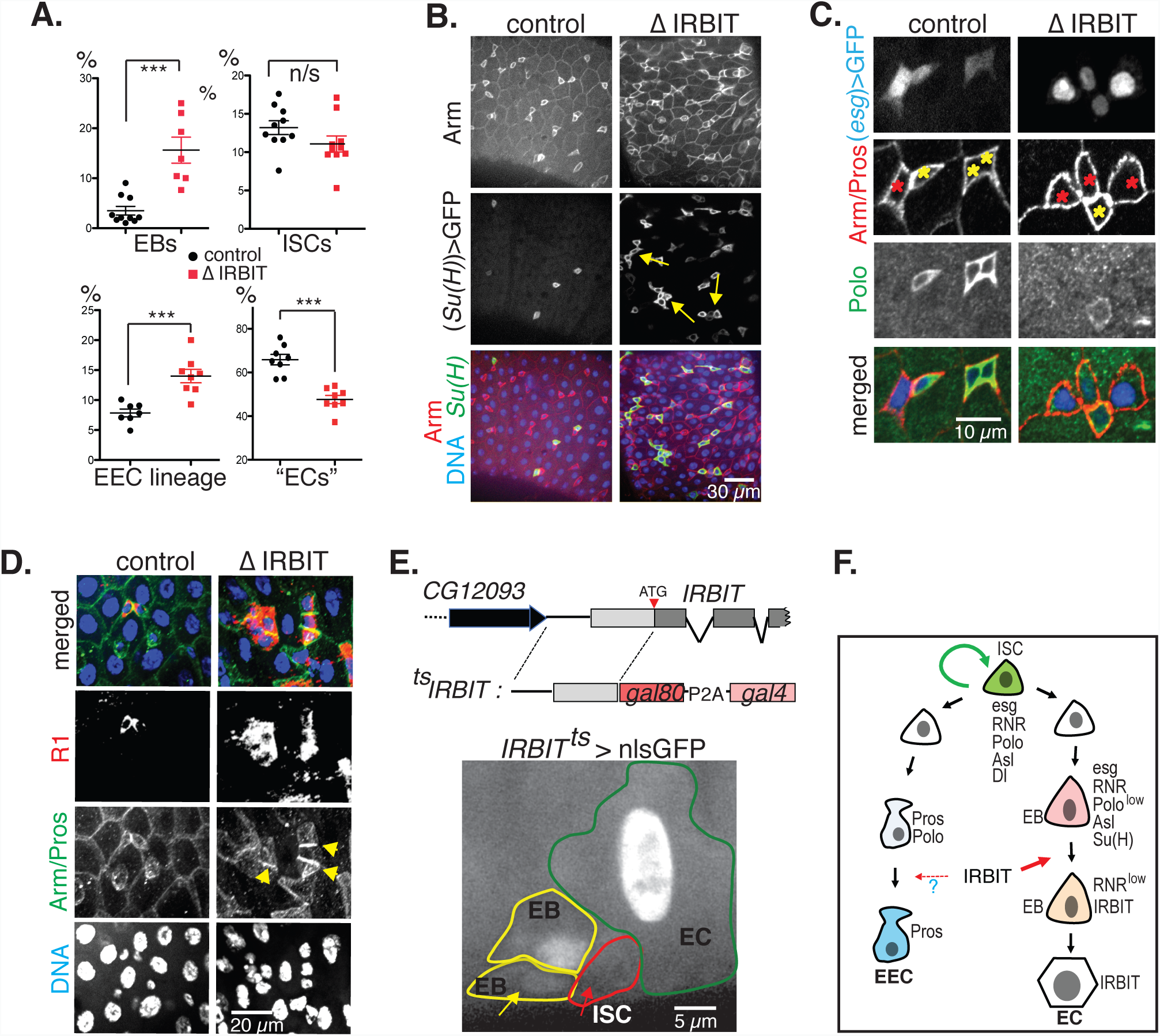
IRBIT mediates differentiation of the ISC progeny. (A) Cell composition in midguts. Control is in black, ΔIRBIT is in red. Quantifications of EBs (Su(H)^+^ cells), ISCs (Delta^+^ cells), cells of EE lineage (Pros^+^ cells) and ECs (large nuclei, Su(H)^−^, Delta^−^, Pros^−^) in pmr of 8 d old female flies. (EECs: N=8 guts; ISCs: N=10 guts; EBs: N=10 (control), N=7 (ΔIRBIT) guts); (ECs: N=10 guts). Error bars represent mean ± SEM. P values derived from unpaired t test with Welch’s correction, n/s, not significant, ***P<0.003. (B) EBs were marked with GFP in 8 d old female guts using temperature-sensitive expression system (^*ts*^*Su(H): Su(H)GBE-Gal4, UAS-mCD8GFP; tub-Gal80*^*ts*^). Note that the cell aggregates in ΔIRBIT (yellow arrows) are GFP^+^. (C) ISC progeny in 8 d old control and ΔIRBIT guts were marked using ^*ts*^esg (*esg-Gal4, UAS-nlsGFP, Gal80^ts^*) (marker of ISCs and EBs, pseudo colored in blue) and probed for Arm (red) and Polo (marker of ISCs, green). Note a single stem cell (high Polo, yellow asterisk) with several (here: 3) attached enteroblasts (GFP^+^, low Polo; red asterisks) in ΔIRBIT. (D) Accumulation of R1^+^ cells in ΔIRBIT. 8 d old female guts stained for Arm (green), R1 (RnrL, large subunit of RNR, red), and DNA (blue). (E) IRBIT is expressed in the ISC progeny. A genomic construct that contains a putative IRBIT promoter and its 5’UTR was fused with *Gal80^ts^-P2A-Gal4* (^*ts*^IRBIT) and used to drive nlsGFP expression (pseudo colored in white). Note the presence of nuclear GFP in EBs (yellow circles) and ECs (green circle) but not in the ISC (red circle, red arrow). Young EB (yellow arrow) is indicated. (F) Summary: IRBIT promotes differentiation of ISC progeny in the EC lineage.

### GATAe stimulates IRBIT expression to suppress RNR and to allow differentiation of the ISC progeny

We next tested whether IRBIT expression is controlled by known transcriptional regulators of ISC progeny differentiation (Zhai et al., 2017). We focused on GATAe, because we noted that during embryogenesis the expression of GATAe (Okumura et al., 2005) is remarkably similar to that of IRBIT. Knockdown of GATAe in Esg-positive cells (ISCs and EBs) reduced IRBIT expression, with concomitant accumulation of undifferentiated progeny showing high R1 expression (Figures 3A, 3B and S5A), suggesting that GATAe is a transcriptional regulator of IRBIT. Although the minimal *IRBIT* promoter (Figure 2E) does not contain “classical” GATAe motif (WGATAR) (Okumura et al., 2005), GATAe stimulates its activity (Figure S5B), indicating that either there is an yet unidentified GATAe motif or/and that GATAe stimulates IRBIT transcription via another transcription factor(s) that lies downstream of GATAe. Importantly, IRBIT overexpression rescued the phenotypic defects in GATAe^RNAi^, indicating that IRBIT is an important downstream target of GATAe in the intestine (Figure 3B). Interestingly, a phospho-mimetic IRBIT mutant that presumably is a much more potent R1 inhibitor (Arnaoutov and Dasso, 2014), rescues GATAe knockdown even better than the IRBIT^wt^ does (Figure S5A), suggesting that phosphorylation of dIRBIT plays an important role during EB maturation.

**Figure 3.**
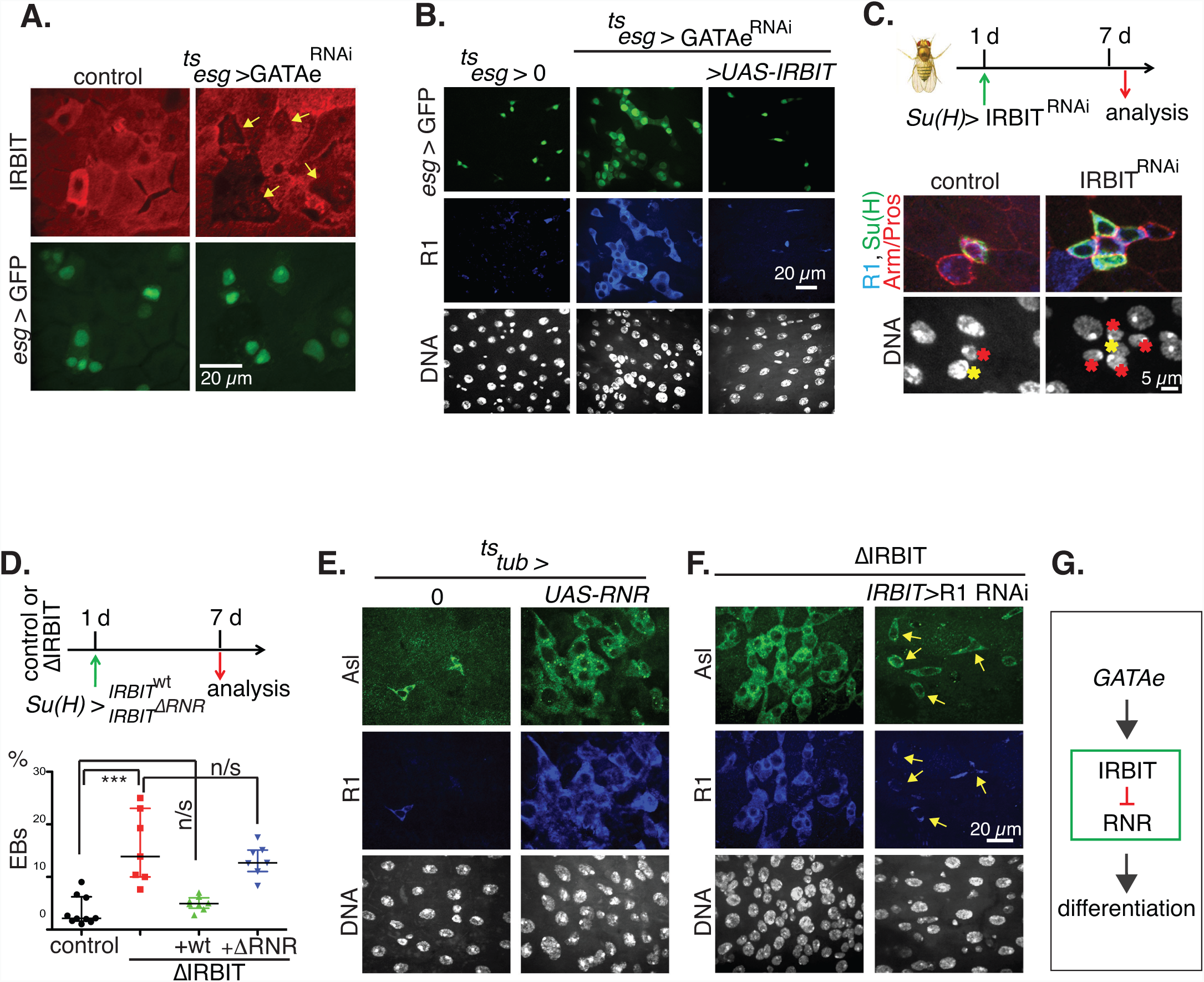
GATAe stimulates IRBIT expression to suppress RNR and to allow differentiation of the ISC progeny. (A) GATAe is required for IRBIT expression in the ISC progeny. The expression of GATAe was silenced with RNAi in the progeny using ^*ts*^esg for 4 d and the midguts were stained with IRBIT antibodies. IRBIT expression of IRBIT was reduced in esg^+^ cells (yellow arrows). (B) IRBIT is a downstream target of GATAe. The expression of IRBIT was induced in GATAe-silenced cells for 7 d (^*ts*^*esg, UAS-GATAe*^*RNAi*^, *UAS-IRBIT*). Note that midguts with reduced GATAe develop RNR^+^/esg^+^ dysplasia, that is rescued by overexpression of IRBIT. (C) IRBIT functions cell-autonomously in EBs. Expression of IRBIT was silenced in EBs (^*ts*^*Su(H), UAS-IRBIT*^*RNAi*^) for 7 d and the accumulation of EBs was monitored by GFP^+^ cells. Note that IRBIT-silenced EBs (red star) fail to detach from mother ISC (yellow star) and accumulate. (D) Quantifications of EBs in pmr of 7 d old ΔIRBIT flies rescued with ^*ts*^*Su(H)>IRBIT* or ^*ts*^*Su(H)>IRBIT*^*ΔRNR*^. N=7-8; error bars represent mean ± SEM. P values derived from the Kruskal-Wallis with Dunn’s multiple comparison test, ***P<0.001. (E) Overexpression of RNR mimics ΔIRBIT phenotype. The expression of RNR (both R1 and R2 (Rnrs)) was induced for 5 d using ^*ts*^*tub* expression system (*tub-Gal4; tub-Gal80*^*ts*^). Note accumulation of progenitor cells (Asl^+^). (F) Suppression of RNR bypasses the requirement for IRBIT in the midguts. The expression of RNR (R1) was silenced in ΔIRBIT midguts by RNAi for 5 d using ^*ts*^*IRBIT* promoter. Note that the ISCs remain intact (yellow arrows). (G) A model of GATAe-IRBIT-RNR pathway.

Genetic manipulations of IRBIT specifically in EBs indicated that IRBIT acts cell autonomously in these cells to mediate their differentiation: IRBIT knockdown induced excessive EBs accumulation in the midgut, whereas IRBIT overexpression in ∆IRBIT EBs restored their normal progression through differentiation (Figures 3C and 3D). Notably, an IRBIT mutant lacking the RNR binding region (aa 53-67, IRBIT^∆RNR^) (Arnaoutov and Dasso, 2014) failed to suppress the EBs accumulation in ∆IRBIT midguts (Figure 3D). In contrast, suppression of RNR activity by hydroxyurea effectively rescued the EBs number and restored the midgut integrity in ∆IRBIT (Figure S5C).

Moreover, overexpression of RNR (using *tub* promoter) resulted in ∆IRBIT-like tissue dysplasia, while silencing RNR using the IRBIT promoter rescued the ∆IRBIT phenotypes (Figures 3E and 3F), strongly indicating that high levels of RNR is detrimental for differentiation and that IRBIT functions to suppress RNR in order to maintain differentiation of ISCs.

Silencing IRBIT expression by using *myo1A* promoter which is expressed in EBs and in ECs recapitulated ∆IRBIT phenotype (Figure S5D). Although we cannot rule out the possibility of additional EC-specific effects of IRBIT that influence EB maturation, our findings collectively indicate that IRBIT is expressed in EBs, and that IRBIT-mediated suppression of RNR in EBs is necessary for ISC progeny differentiation (Figure 3G). Consistent with this conclusion, RNA-Seq analysis of midguts further indicated that IRBIT promotes differentiation of ISC progeny (Figure S6; Table S1). Importantly, accumulation of undifferentiated progeny in ∆IRBIT midguts was not dependent on intestinal bacterial load *per se*, because we observed similar phenotype in axenic flies, indicating that the problem of ISC differentiation was not a result of microorganism-induced inflammation (Figure S6A; Table S1).

Of note, we observed a similar distribution of IRBIT in mouse jejunum (middle part of the small intestine), with high levels in differentiated ECs but low levels in the ISC niche (Haber et al., 2017) (Figure S7), suggesting that IRBIT may play a similar role in mammalian ISC differentiation.

### Maintenance of IRBIT-RNR regulatory circuit prevents formation of age-related phenotype in the intestine

The characteristic tissue dysplasia that rapidly develops in ∆IRBIT or in GATAe^RNAi^ midguts is reminiscent of dysplasia in aging flies and mammals (Jasper, 2015), prompting us to ask whether loss of the GATAe-IRBIT-RNR pathway may underlie loss of intestinal homeostasis with age. To address this, we performed RNA-Seq analysis of midguts under a variety of conditions. We measured expression genome-wide and assessed significant differential expression (DE) in pairwise comparisons. To visualize DE, we performed a hierarchical clustering analysis of log ratios of DE genes. Changes in gene expression pattern during normal intestinal aging were opposite to the changes induced by IRBIT expression in young guts (Figures 4A and S8A; Table S2), suggesting that IRBIT protects guts from age-induced changes in transcription. We also analyzed the anti-microbial response (AMR), an innate immune mechanism that helps flies control intestinal microbiota. The AMR increases with age because the frail aging epithelium becomes more susceptible to bacterial infection (Broderick et al., 2014; Chen et al., 2014; Regan et al., 2016). Gene expression changes in young ∆IRBIT midguts showed induction of genes associated with the AMR, consistent with the idea that these midguts had prematurely developed characteristics similar to age-associated frailty (Figures 4B, S8B). In addition, we also observed a significant overlap between genes those regulation is controlled by both IRBIT and Sox21a (Chen et al., 2016; Zhai et al., 2017), suggesting that ∆IRBIT midguts might accumulate EBs at their Sox21a-dependent stage of differentiation (Figure S8C; Table S3). Because GATAe may act downstream of Sox21a (Zhai et al., 2017), this data supports our model in which IRBIT is an effector of GATAe.

**Figure 4.**
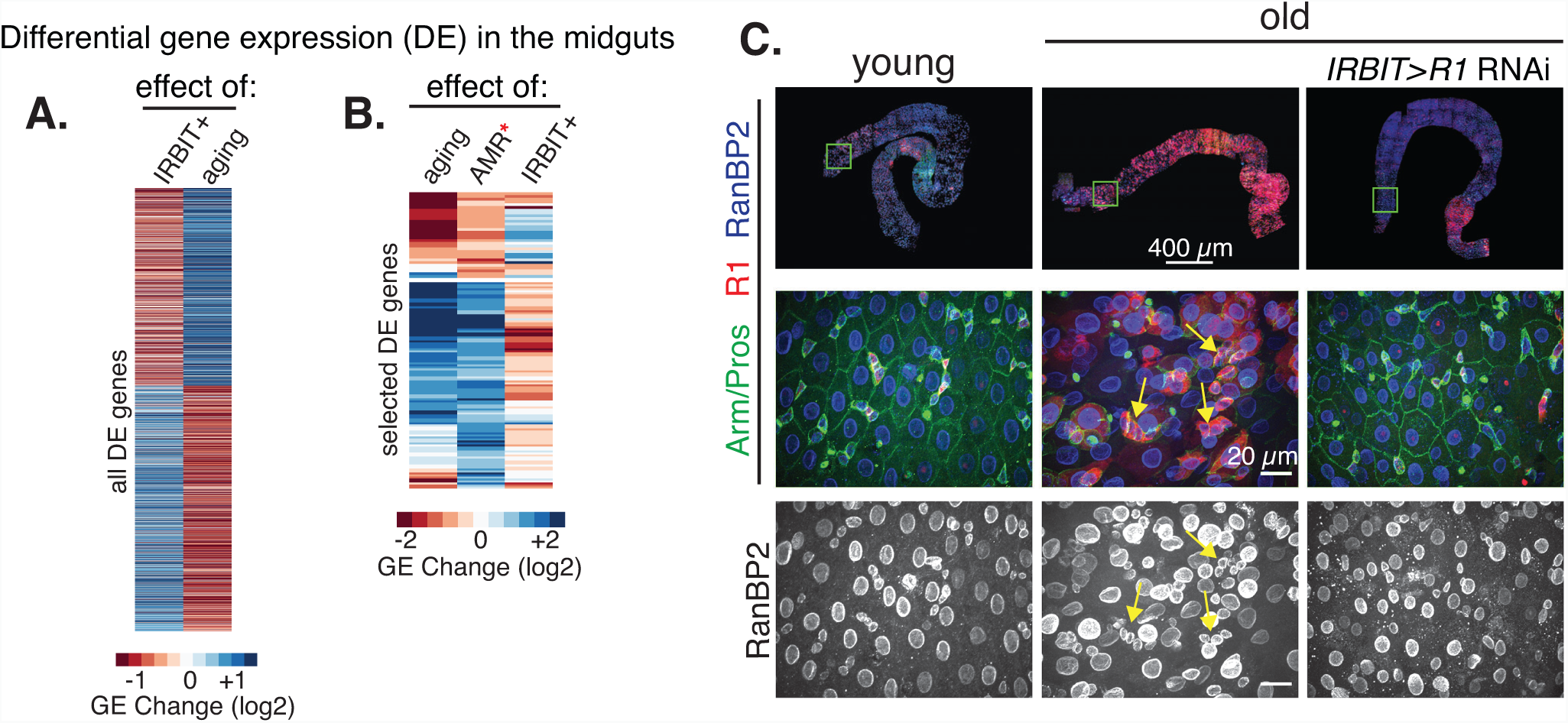
IRBIT is required for intestinal homeostasis. (A) The loss of IRBIT mirrors aging program in the gut. Clustering of “IRBIT+” (comparison of gene expression in 8 d old midguts control (*yw*) vs. ΔIRBIT) and “aging” (comparison of gene expression between *yw*, 40 d old and *yw*, 8 d old) DE genes. Note the strong anticorrelation of DE between “aging” and “IRBIT+” (Pearson’s r = – 0.54, p < 2.2e-16, F test). (B) ΔIRBIT midguts elicit strong AMR. Unsupervised hierarchical clustering of AMR genes (Broderick et al., 2014)* with “IRBIT+” and “aging” -dependent genes. We performed separate hierarchical clustering for those upregulated genes (the top) as well as downregulated genes (the bottom). Note the anti-correlation of “IRBIT+” and the AMR response. (C) Maintenance of IRBIT-RNR pathway prevents formation of aging phenotype in the intestine. RNR (R1) was continuously silenced in midguts by RNAi using ^*ts*^IRBIT driver and the midguts of 40 d old flies were stained with Arm/Pros (green), R1 (red) and RanBP2 (nuclear pores, blue). Note the disappearance of dysplasia (yellow arrows) and the maintenance of normal tissue architecture.

Moreover, midguts of old flies develop RNR-positive clusters of undifferentiated cells, similar to young ∆IRBIT or GATAe^RNAi^ midguts (Figure 4C). Importantly, reducing RNR levels in the ISC progeny but not in the ISCs by using the *IRBIT* promoter antagonized dysplasia and restored gut tissue integrity, suggesting that the GATAe-IRBIT-RNR pathway plays an important role during intestinal homeostasis.

## Discussion

IRBIT acts as an allosteric inhibitor of RNR (Arnaoutov and Dasso, 2014). Here, we have examined the biological importance of this mechanism using the midgut of *Drosophila*, a well-established system for stem cell function and differentiation. Collectively, our results indicate that IRBIT is required for maturation of EBs into ECs, and our data suggest that IRBIT acts through RNR in this process. Numerous findings lead us to this interpretation: First, IRBIT expression correlates with the EB maturation (Figure 2E). Second, IRBIT loss-of-function resulted in accumulation of EBs (R1^+^, Su(H)^+^ cells) (Figures 2 and 3). Third, suppression of IRBIT specifically in EB mirrored ∆IRBIT phenotype (Figure 3C). Fourth, suppression of RNR activity in ISC progeny that stalled in ∆IRBIT midguts promoted their differentiation (Figure 3F), and fifth, expression of IRBIT^wt^ but not IRBIT^∆RNR^ specifically in EBs rescued the ∆IRBIT phenotype (Figure 3D). Based on these observations, we propose a model in which IRBIT acts in newly formed EBs to attenuate RNR and to induce differentiation. As RNR is expressed in both ISC and EB, the absence of IRBIT inhibition of RNR activity presumably results in an ISC-level dNTP pool within EBs and delays their differentiation. Because the differentiation of EBs is critical for replenishing aging ECs, the block of differentiation in the absence of IRBIT ultimately results in frail midgut epithelium that cannot maintain strong anti-bacterial protective wall (Figures S2C and S2D), causing continuous immune response (Figure 4B).

Phenotypically, IRBIT deletion (with an accumulation of undifferentiated EBs without accumulation of ISCs) is similar to the phenotypes that had been prior observed in flies where expression of Sox21a, GATAe, JAK/STAT or Dpp were silenced (Zhai et al., 2017). Our results suggest that IRBIT acts downstream of GATAe, because suppression of GATAe expression in ISC progeny resulted in accumulation of progeny with reduced IRBIT levels. Moreover, the progeny in GATAe^RNAi^ midguts, as in the case of ∆IRBIT midguts, remained positive for R1, indicating an incomplete suppression of RNR in these cells. In addition, overexpression of IRBIT stimulated differentiation of EBs, stalled in the absence of GATAe, indicating that IRBIT is an important downstream target of this transcription factor.

The C-terminal domain of IRBIT shares significant homology to S-adenosyl homocysteine (SAH, AdoHcy) hydrolase (SAHH), a crucial enzyme that is responsible for the removal of a byproduct of the methylation reaction. It has been suggested that IRBIT acts as a negative regulator of canonical SAHH enzyme to regulate methionine metabolism in flies (Parkhitko et al., 2016). We have performed extensive biochemical and genetic experiments to test whether this domain could work either as a SAHH or as a natural inhibitor of SAHH. The full-length IRBIT or IRBIT’s core domain did not show SAHH activity, binding to canonical SAHH nor the capacity to interact with AdoHcy agarose, indicating that IRBIT does not bind SAHH or the SAHH substrate (Figure S9). Therefore, we consider it unlikely that IRBIT acts as a dominant negative form of SAHH (Parkhitko et al., 2016). Our biochemical, genetics and histology results strongly indicate that IRBIT’s N-terminus is critical for its function as an inhibitor of RNR (Arnaoutov and Dasso, 2014). We speculate that the function of the IRBIT’s core domain is to form a dimer interface between the two IRBIT molecules for proper positioning of their intrinsically disordered N-termini, thus allowing them to interact with IRBIT-corresponding partners that, too, typically exist as dimers, or as a complex of dimers, like RNR (Arnaoutov and Dasso, 2014).

As suppression of the cell cycle promotes differentiation, the choice between proliferation and differentiation may, in principle, be controlled by many cell cycle checkpoint components (Ruijtenberg and van den Heuvel, 2016). Our results indicate that the maintenance of dNTP levels could be one such mechanism. The maturation of EBs could be viewed as a two-step process: first, Notch-mediated signals pause the ISC daughter in G1 and induce a strong Su(H) expression to commit it to the EB lineage; second, a committed EB undergoes polyploidization to fully mature into an adult EC. We initially suspected that IRBIT-mediated control of RNR in EBs might be essential for their polyploidization, i.e. switching replication/mitotic program into endoreplication. Based on our data in IRBIT-depleted HeLa cells (Arnaoutov and Dasso, 2014), we reasoned that the general speed of the replication fork progression could be the trigger point behind such mechanism, and the delay in endoreplication would result in the accumulation of EBs that are not fully endoreplicated. We tested this hypothesis by an artificial reduction of RNR activity in EBs that had been accumulated in ∆IRBIT guts. To our surprise, the administration of HU at 20 mM to the diet of ∆IRBIT flies that already accumulated undifferentiated progeny rapidly reduced the amount of EBs. More importantly, suppression of RNR abundance in ISC progeny using RNAi, driven by *IRBIT* promoter also rescued tissue dysplasia in ∆IRBIT midguts. Although it is formally possible that inhibition of RNR may have resulted in clearance of progenitors by apoptosis or by some other mechanism, we favor the idea that it is the presence of high RNR activity in EBs that is detrimental for the initial step of EBs maturation and that the decrease of the RNR activity is necessary for the decrease of Su(H) signal and normal progression of EBs into ECs, even if endoreplication is not completed. We speculate that RNR activity within the newly formed EBs must be suppressed by IRBIT to a certain threshold in order to pause them in early S phase and to proceed with their differentiation. Once the EB is fully committed to this transition, it commences endoreplication, that also could be under the control of IRBIT, consuming endogenous dNTP produced by the residual activity of RNR. Alternatively, endoreplication of ECs may rely upon deoxynucleosides that could be absorbed from the gut lumen.

In summary, we have shown a novel role of IRBIT and RNR during homeostasis of midgut epithelium. IRBIT expresses in postmitotic intestinal stem cell progenitors to suppress RNR and to assist their differentiation into adult epithelial cells, a process that is essential for sustainability of the tissue during animal’s lifespan. Our study provides a novel approach toward understanding dysplasia, potentially facilitating development of strategies that could help containing intestinal diseases.

## Materials and Methods

Details of materials and methods including fly husbandry, genotypes, antibodies, plasmids and assays are described in Supplemental Information, Materials and Methods.

## Supporting information

Reagent table

## Acknowledgements

This study was supported by 2018 NICHD DIR Director’s Award. We are grateful to K. Ten Hagen and N. Rusan for reagents. We are also indebted to M. Jaime, L. Fu and YB Shi for their help and technical assistance. We thank M. Lilly and O. Demidov for the critical reading of the manuscript. We also thank the Bloomington Stock Center at the University of Indiana and VDRC for fly stocks, and the Developmental Studies Hybridoma Bank at the University of Iowa for antibodies.

## Materials and Methods

### Fly stocks

*IRBIT^70^* and *IRBIT^130^* were obtained by imprecise excisions of a transposable element, *P[EP]AhcyL1[G4143]* (BL-27147), inserted in the 5’UTR of *IRBIT* gene. The resulting lines were characterized by genomic PCR and DNA sequencing. The genomic fragments removed were as follows: *IRBIT^70^*, 3L: 2770370-2772267 (genome release 6), including exon 1 and part of exon 2; *IRBIT^130^*, 3L: 2770256-2772267 (genome release 6), including exon 1 and part of exon 2 of the predicted *IRBIT* gene. The heteroallelic combination *IRBIT^70^/IRBIT^130^* produced viable and fertile flies (referred to as ∆IRBIT in this study). *UAS-IRBIT* transgenic lines were generated by insertion of *IRBIT* cDNA or mutant variants in a pUAST vector containing an attB sequence followed by phiC31 integrase-mediated germline transformation at VK00018 docking site at cytological location 53B2 on the second chromosome (BL-9736). The genomic fragment, corresponding to III: 2768343-2772511 and comprising the IRBIT gene and the flanking regions, was also introduced at this docking site (BL-9736). Introduction of this fragment into ∆IRBIT flies produced a viable stock that we refer to as ∆IRBIT^Resc^. A genomic construct that contains a putative IRBIT promoter and its 5’UTR was fused with *GAL80^ts^-P2A-GAL4* to generate ^*ts*^IRBIT line. *UAS-RNR* transgenic line was generated by insertion of *R1 (rnrL) and R2 (rnrS)* cDNAs, linked with P2A sequence into a pUAST vector followed by integration at VK00018 docking site, as described above. Transgenic animals were generated by BestGene, Inc. using standard phiC31 integrase-mediated germline transformation protocols.

For survival analysis, 40 virgin flies of each genotype were collected. Flies were reared on standard Bloomington food and kept at 25^0^ C on a 12 hour light/dark cycle. Flies were flipped onto new food 1-2 times per week and dead flies were counted until there were no survivors.

### Preparation of axenic flies

Axenic flies were prepared according to (Broderick et al., 2014) with modifications. Briefly, 40-80 flies were kept overnight in collection cages with freshly prepared agar juice plates (Genesee, Inc). Embryos (0-16h old) were collected, washed several times with PBS using a fine mesh and transferred to 9-well glass depression plates (Corning). PBS was aspirated and the embryos were rinsed with 70 % ethanol followed by incubation with 4% sodium hypochlorite for 10 min at RT. Embryos were then washed 3 times in cell culture grade water and transferred to a 6 cm dish, containing axenic food (Bloomington formulation (Genesee, Inc)), autoclaved, cooled to 50^0^ C and supplemented with 1x Pen/Strep mixture (Life Technologies)). After hatching, the flies were transferred to standard vials containing axenic food. The axenic status of these animals was confirmed by plating 3 homogenized flies onto LB plate. No colonies were detected after 3 d incubation at RT. The absence of RNA transcripts corresponding to anti-bacterial peptides (DptA, DptB, AttC) in RNA-Seq data serves as an additional control for the verification of axenicity.

### Preparation of midguts for RNAseq

For the initial screen (as in Figures 4A and 4B) the flies were reared on Jazz-Mix food (Fisher Scientific). For each genotype, 30 freshly eclosed females (1-10 hours post eclosion) were transferred and reared for the indicated time into a new food vial supplemented with 100 µl of autoclaved yeast paste (20% v/v). Flies were then anesthetized and dissected in PBS. 5-6 midguts (a region between Malpighian tubule juncture and cardia) were collected into a single eppendorf tube containing 1 ml of Trizol. Care was taken to ensure that the whole procedure took no more than 20 min. Total RNA was isolated according to the manufacturer’s instructions and stored at −80° C. Three independent replicas for each analysis were prepared. 2 µg of each total sample RNA, mixed with 10 pg of control Spike-in RNAs (NIST) were processed to create cDNA libraries using TruSeq stranded mRNA kit (Illumina).

For the follow-up screen and for most of the analysis, the flies were reared on Bloomington food alone (for normal conditions), or containing 1x penicillin-streptomycin mixture (axenic conditions). 20 mM HU was added to the autoclaved and cooled (~50°C) food, where needed. The guts were dissected and processed as above.

RNA-Seq libraries were sequenced on the Illumina platform (HiSeq 2500, Illumina, San Diego, CA) at NIDDK Genomics Core (Bethesda, MD) as 76 bp read-lengths and single-ended. CASAVA 1.8.2 (Illumina) was used for base-calling. We mapped RNA-Seq reads onto Drosophila reference genome Release 6 (Hoskins et al., 2015) (major scaffolds only: chromosomes 2, 3, 4, X, Y, and mitochondrial genome) with ERCC Spike-in RNA sequences (Zook et al., 2012). We used TopHat 2.1.1 (Kim et al., 2013) for the mapping with parameters -g 1 and -G. For the latter, we provided FlyBase gene annotation model 6.12 (Marygold et al., 2016). From the mapping results, we used HTSeq 0.6.1p1 (Anders et al., 2015) to obtain gene-level read counts. We used “-s reverse” parameter to correspond strandness but otherwise with the default setting. We calculated gene-level FPKM values (Fragments Per Kilobase of transcript per Million mapped reads) based on each gene’s maximum transcript length from the collapsed gene model. We used RSeQC (Wang et al., 2012) to inspect RNA quality based on the coverage uniformity over the gene body. We measured ratios of the mean coverage between 20 to 40 percentiles of scale gene bodies over that between 60 to 80 percentiles. Based on the ratio, we discarded the samples that display 3’-biased coverage (e.g. the ratio < 0.9), which have potential RNA degradation, from our downstream analysis (total 5 samples). All remained replicates have FPKM correlations greater than 0.95 (Pearson’s r). We filtered out non-protein coding genes before differential expression analysis. Also, we used expression cutoffs based on intergenic FPKM signals as described previously (Lee et al., 2016; Zhang et al., 2010). We determined FPKM signals based on read counts and intergenic lengths between two annotated gene models that are adjacent each other. We set top 95 percentile of such signal (FPKM 1.90 and 1.43 for non-axenic and axenic samples, respectively) as our expression cutoff with estimated false discovery rate at 5%. We measured differential expression of genes that are over the cutoff from at least 5% of the total samples for each sex, and also that have more than 1 CPM (Counts per Million mapped reads) from any of the all samples. We used DESeq2 1.14.1 (Love et al., 2014) for our differential expression analysis. DE genes are whose change is significant at p values corrected with Benjamini and Hochberg method < 0.05. Short read sequences as well as gene expression levels in counts and FPKM values are available in Gene Expression Omnibus (Edgar et al., 2002) with an accession ID of GSE109862 (token: wvqzoasqdnghxef).

### Antibody generation and Immunofluorescence

The following antibodies against *Drosophila* proteins were generated: IRBIT – raised in rabbits against baculovirus-expressed 6His-tagged fragment of dIRBIT (aa 1-106); R1 – raised in rabbits against baculovirus-expressed full length 6His-tagged RnrL; RanBP2 – raised in chickens against dRanBP2 (aa 2318-2696); Polo – raised in rabbits against bacterially-expressed full length 6His-tagged Polo. Antibodies were developed in Pacific Immunology (Ramona, CA) and then affinity-purified using a GST-fusion of a corresponding antigen.

For immunohistochemical analyses, 3-10 whole guts per genotype were dissected in PBS, transferred to a siliconized Eppendorf tube containing 4 % paraformaldehyde in PBS, and fixed for 1 h at RT on a rotator. The samples were pelleted at 500 g for 10 s, washed three times for 10 min with TBS-T, containing 1 % TX-100, then blocked for 1 h at RT in TBS-T, containing 10 % normal goat serum. The samples were incubated overnight at 4 C with the primary antibodies, then pelleted (500 g for 10 s) and washed 3 times 20 min each in TBS-T and incubated with the corresponding secondary antibodies (1:500; Alexa-labeled goat anti-rabbit, anti-mouse, anti-chicken, ant-guinea pig (Invitrogen)) and Hoechst 33342 for 2 h at RT on a rotator. The specimen were washed as above, then mounted in ProLong Diamond Antifade Mountant (ThermoFisher Scientific). To analyze the peritrophic membranes, 3-4 whole guts per genotype were fixed in 1 ml of 2 % glutaraldehyde/PBS for 1 h on a rotator. The guts were pelleted as above, washed twice with blocking buffer (TBS-T, containing 100 mM TrisHCl (pH 8.0), 30 mM glycine HCl (pH 5.0) and 1 % TX-100) and transferred in 50 µl of the blocking solution to a well of the 9-well glass depression plate. Posterior midgut region was torn by forceps and microneedles and the peritrophic membrane protrusion were visualized. The buffer was carefully aspirated and 200 µl of staining solution (TBS-T, containing 10 % goat serum and lectin HPA, labelled with Alexa Fluor488 (1:500)) was added to the dissected guts and the plate was incubated for 2 h on a rocket shaker. The guts were then mounted in ProLong Diamond Antifade Mountant with DAPI.

The following primary antibodies were used: IRBIT (1:300); R1 (1:100), RanBP2 (1:300), Polo (1:100), Armadillo (1:100, DSHB), Asl (1:1000) (Rusan and Peifer, 2007), Prospero (1:100, DSHB), Discs large (1:200, DSHB), Delta (1:100, DSHB), Actin (1:500, Sigma), Tubulin (1:500, clone DM1A, Sigma).

### Transmission electron microscopy

Midguts of adult flies were dissected in PBS, fixed in 2.5% glutaraldehyde/2% formaldehyde/2 mM CaCl2 in 0.1 M cacodylate buffer pH 7.4 for 30 minutes at room temperature followed by 1.5 hour on ice in fresh fixative. After 5 washes in the buffer, the samples were postfixed in 2% osmium tetroxide in the same buffer for 1.5 hours on ice, washed once in the buffer and 5 times in double distilled water. The samples were then stained en bloc overnight in 2% aqueous uranyl acetate, washed twice in water and dehydrated in series of ethanol concentrations (30%, 50%, 70%, 90%, 3×100%), and further penetrated with EMbed 812 epoxy resin (EMS, Hatfield, PA) diluted 1:2, 1:1 (1 hour each) and 2:1 (overnight, open vial) in propylene oxide, followed by incubation twice in undiluted fresh resin (2 hours each). The midguts were embedded in flat molds and polymerized for 60 hours at 65°C. Thin sections (70-80 nm) were cut on Leica Ultracut UC7 microtome (Leica, Deerfield, IL), mounted on formvar-carbon coated grids and stained with uranyl acetate. The samples were examined on FEI Tecnai 20 TEM (FEI, Hillsboro OR) operated at 120 kV and images were recorded on AMT XR81 CCD camera (AMT, Woburn, MA).

### RNA FISH

Mouse small intestine (from 4-8 week old adult mice) was fixed in 4% pfa for 24 h and embedded in paraffin; 5 µM sections of jejunum were cut and placed on round coverslips followed by paraffin removal, dehydration and rehydration in ethanol series using standard techniques. Coverslips were then incubated in TE buffer for 10 min at 95^0^ C, followed by incubation in protease solution (4 µg Proteinase K in PBS) for 10 min at RT. Tissue sections were then washed 3 times with PBS-T and post-fixed in 4% pfa in PBS for 1 h, washed twice in TBS-T and processed for single molecule RNA FISH using ViewRNA Cell Plus Assay kit (ThermoFisher).

### Biochemical experiments

Recombinant 6His-hSAHH, 6His-hIRBIT, GST-hIRBIT, GST-hIRBIT2, GST-dIRBIT, GST-dCG8956 (IRBIT-like) and dR1 proteins were prepared in baculovirus-expressed Sf9 insect cells as described (Arnaoutov and Dasso, 2014). dATP-dependent association of R1 with IRBIT proteins were performed as described (Arnaoutov and Dasso, 2014). For WB analysis (Figure 1B), three female flies of a corresponding genotype were homogenized in 200 µl of PBS, mixed with 200 µl of 5 × SDS-PAGE sample buffer and boiled for 10 min. Samples were vortexed and centrifuged at 250000 g for 10 min at RT on an ultracentrifuge (Beckman). Supernatant were run on SDS-PAGE and probed with antibodies against IRBIT and Tubulin. For SAH hydrolysis assay (conversion of S-adenosylhomocysteine (AdoHcy) into adenosine and homocysteine), 100 ng of a corresponding recombinant protein or 20 mkl of immunoprecipitated IRBIT was added to the 100 µl reaction buffer containing 20 mM TrisHCl (pH 8.0), 100 mM NaCl, 1 mM DTT, 1 µM NAD, 1 mM AdoHcy, 0.1 unit adenosine deaminase and incubated for 15 min at RT. The conversion of adenosine to inosine was monitored on NanoDrop. For estimation of the reverse reaction (synthesis of AdoHcy from adenosine and cysteine), recombinant or immunoprecipitated proteins were added to the 100 µl reaction buffer containing 20 mM TrisHCl (pH 8.0), 100 mM NaCl, 1 mM DTT, 1 µM NAD, 1 mM adenosine, 1 mM homocysteine, and incubated for 15 min at RT. Adenosine deaminase (1 unit) was added either before or after the completion of the reaction. The abundance of adenosine peak was monitored using NanoDrop.

For pulldown experiments, AdoHcy was immobilized on agarose using 2,4,6-trichloro-1,3,5-triazine. 20 µl of AdoHcy-agarose was mixed with recombinant hIRBIT and SAHH (1 µg each) in 600 µl of binding buffer (20 mM TrisHCl (pH 8.0), 100 mM KCl, 1mM DTT), containing 1 mg/ml BSA and incubated for 1 h at 4^0^ C on a rotator. The beads were pelleted (400 g, 10 s) and washed twice with 1 ml of binding buffer. Bound proteins were eluted by boiling the beads in 50 µl of SDS-PAGE sample buffer, and Western blot was probed with IRBIT and SAHH antibodies.

### Yeast strain constructions

Heterozygous diploid yeast strain 20176 (SAH1, YER043C) was transformed with SAH1 Ura^+^ plasmid. The Ura^+^ transformants were sporulated and subjected to tetrad analysis. An Ura^+^ ascospore clone that was incapable of growth on medium containing 5-fluoroorotic acid (5-FOA) was picked and transformed with *LEU2* plasmid containing either scSAH1, hSAHH, hIRBIT or hIRBIT domain (aa 105-530), that has high homology to SAHH. Cells were plated on synthetic media either lacking Ura and Leu or on media, lacking Leu and containing 5FOA. The growth on Leu/5FOA plate would indicate successful rescue of SAH1 deletion.

---

**Figure S1.**
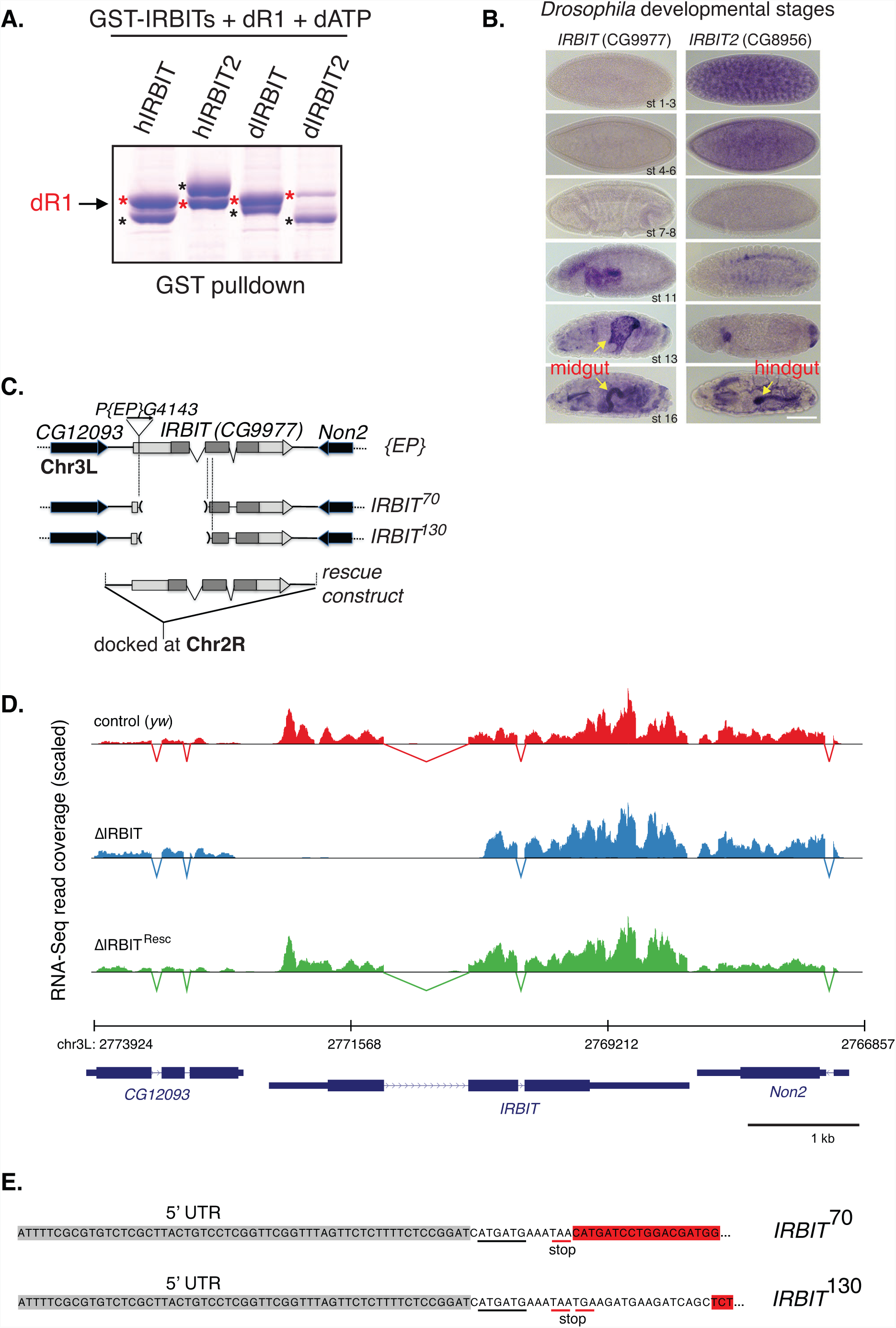
Characterization of *Drosophila* IRBIT. (A) Interaction of IRBIT and IRBIT2 with RNR. 5 µg of purified GST-tagged hIRBIT, hIRBIT2, dIRBIT and dIRBIT2 were incubated with 50 µg recombinant *Drosophila* R1 in the presence of 10 µM dATP and the complexes were precipitated by Glutathione-agarose. Note that in all cases, except for dIRBIT2 there is a strong dATP-mediated interactions between R1 (Red asterisks) and IRBITs (blue asterisks). (B) Expression of IRBIT and IRBIT2 mRNA during embryogenesis. Note the non-overlapping pattern of expression. Midgut and hindgut are indicated with arrows. (C) Schematics of the *IRBIT* locus and the *P[EP]G4143* transposable element, the small lesions corresponding to the IRBIT alleles and the genomic rescue construct. (D) RNA-Seq read coverage of the *IRBIT* locus in the *yw*, ∆IRBIT and ∆IRBIT^Resc^ midgut samples. (E) ∆IRBIT does not code a functional protein. RT-PCR analysis of two null IRBIT alleles. IRBIT mRNAs coded by *IRBIT^70^* and *IRBIT^130^* alleles. The grey region denotes 5’UTR and the red region denotes the remaining of the second exon. Note that the fragment (white region), left by the excised transposable element includes two putative initiating codons (underlined), followed by an immediate stop codon.

**Figure S2.**
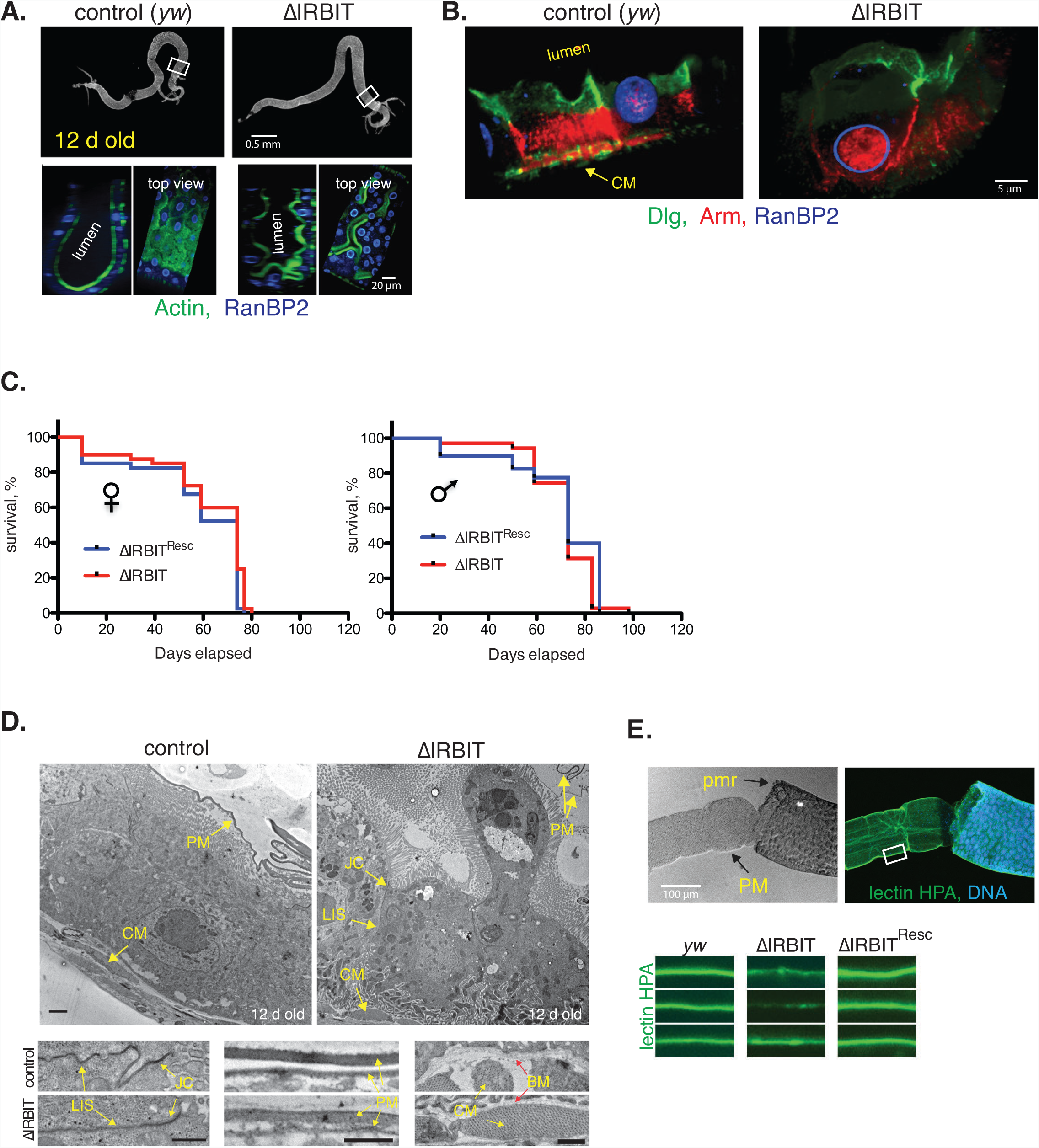
∆IRBIT mutant midguts have aberrant organization. (A) Related to Figure 1C. Midguts of 12 d old control and ∆IRBIT flies (same as in Figure 1C) were stained with antibodies against actin and RanBP2 (nuclear pores), and the pmr (white box) was reconstructed by 3D analysis using Volocity package. Note multiple epithelial polyps protruding in the lumen of ∆IRBIT midguts. (B) Midguts of 12 d old control (Hoskins et al.) and ∆IRBIT flies stained with antibodies against Dlg (Discs large, detects tight junctions), Arm (Armadillo, detects adherence junctions), and RanBP2 (nuclear pores). Two neighboring enterocytes (EC) in each genotype was reconstructed by 3D analysis using Volocity package. Note both aberrant EC morphology and a weak Arm staining between ECs in ∆IRBIT. (C) Lifespan of Drosophila does not depend on IRBIT. 40 virgin flies of indicated genotype and sex were collected. Flies were reared on standard Bloomington food and kept at 25^0^ C on a 12 hour light/dark cycle. Flies were flipped onto new food 1-2 times per week and dead flies were counted until there were no survivors. The change of lifespan between ∆IRBIT and ∆IRBIT^Resc^ flies was not significant (as assayed by both Mantel-Cox and Gehan-Breslow-Wilcoxon tests). (D) Posterior midguts of 12 d old *yw* and ∆IRBIT flies were analyzed by electron microscopy. BM, basement membrane; CC, cell contacts; CM, circular muscles; PM, peritrophic membrane; JC, junctional complex (tight junctions), LIS, lateral intracellular septum (adherence junctions). Magnified regions that cover BM, CC and PM are shown. Note the less dense structure of PM and LIS in ∆IRBIT samples. (E) IRBIT controls maintenance of peritrophic membrane (PM). Top: Pmr of 8 d old axenic *yw* flies was fixed, detached and stained with lectin-HPA and DAPI to visualize PM and DNA, respectively. The phase contrast and the fluorescent image of the same region is shown. Bottom: PMs of 8 d old axenic *yw*, ∆IRBIT and ∆IRBIT^Resc^ flies. Note the decrease in intensity of lectin-HPA-positive signal in the ∆IRBIT pmr.

**Figure S3.**
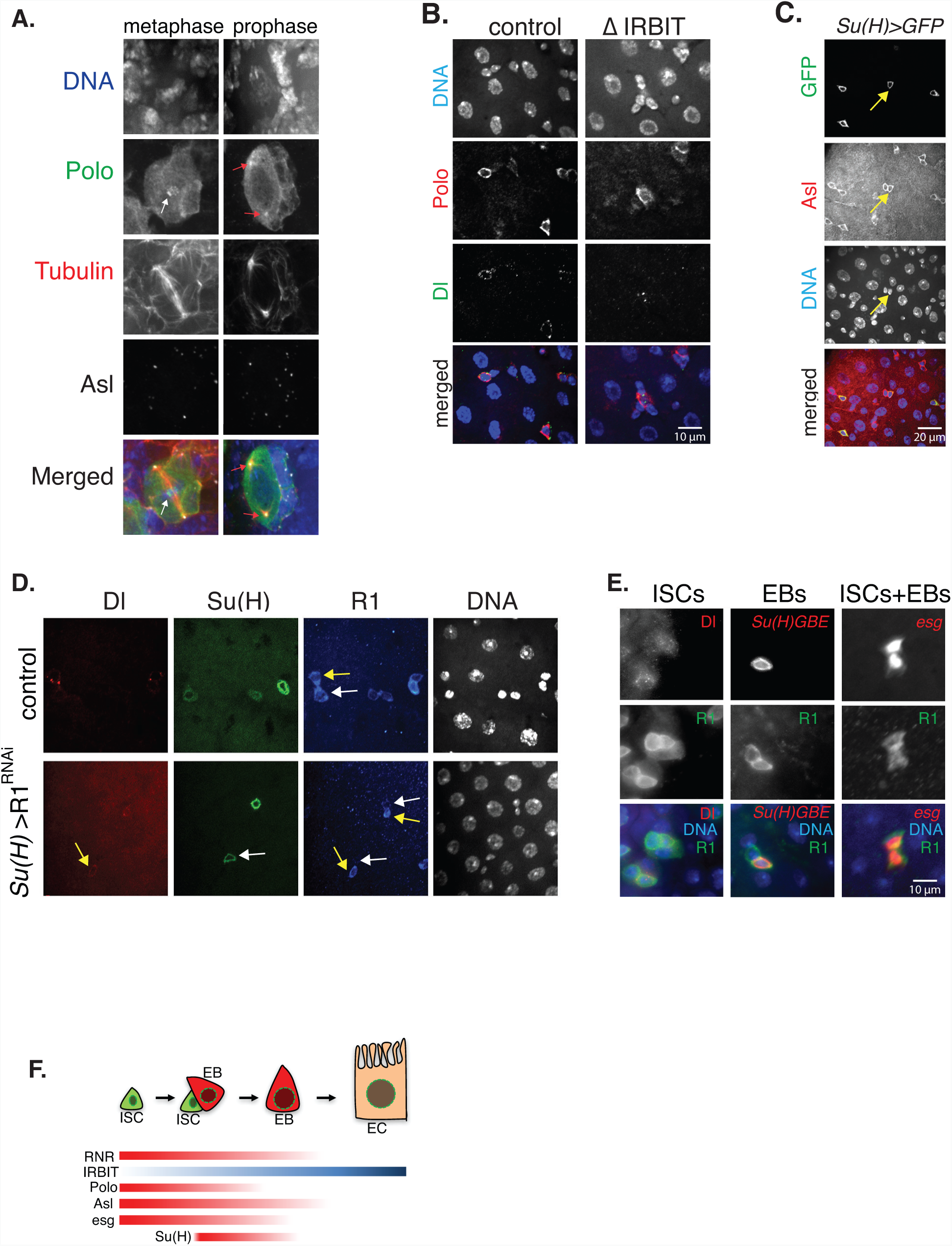
Characterization of ISC- and EB-specific markers. (A) Specificity of Polo antibodies. Third instar larvae brains of *yw* flies were dissected, fixed and stained with antibodies against Polo, Asterless (Asl) and Tubulin. DNA was counterstained with DAPI. Arrows indicate the position of MTOC (microtubule-organizing center; Asl-positive, red arrows) and kinetochores (white arrow) during neuroblasts’ anaphase and metaphase, respectively. (B) Midguts of 8 d old *yw* and ∆IRBIT flies were stained with antibodies against Dl (Delta, ISC marker) and Polo. DNA was counterstained with Hoechst 33342. Note that Polo is highly expressed in the ISC and, to a lower degree, in EBs. (C) Midguts of 8 d old flies, labelled with a temperature sensitive trinary expression system: ^*ts*^*Su(H): Su(H)GBE-Gal4, UAS-GFP.mCD8; tub-Gal80ts* (that specifically labels EBs) and co-stained with antibodies against Asterless (Asl, red). DNA was counterstained with Hoechst 33342 (blue). Note that both ISC and EB express Asl at the similar levels. The yellow arrow denotes an ISC-EB pair. (D) Specificity of R1 (RnrL, large subunit of RNR) antibodies. The midguts of 1 d old flies were silenced with R1 RNAi for 5 days using temperature sensitive system: ^*ts*^*Su(H); UAS-R1*^*RNAi*^ that allows induction of R1^RNAi^ specifically in EBs. Guts were co-stained with antibodies against Dl (red) and R1 (blue). Note the reduction of R1 signal in the EB (Su(H)^+^ cells), but not in ISC (Dl^+^) cells. (E) Midguts of 8 d old flies marked with either ^*ts*^esg>GFP (allows GFP expression specifically in the ISCs and EBs) or ^*ts*^Su(H)>GFP (allows GFP expression specifically in EBs) were co-stained with antibodies against Dl and R1. Note that both ISC and EB abundantly express R1. (F) Schematics of differential expression of RNR (R1), IRBIT, Polo, Asl with regards to Esg and Su(H) markers during the ISC-EB-EC transition.

**Figure S4.**
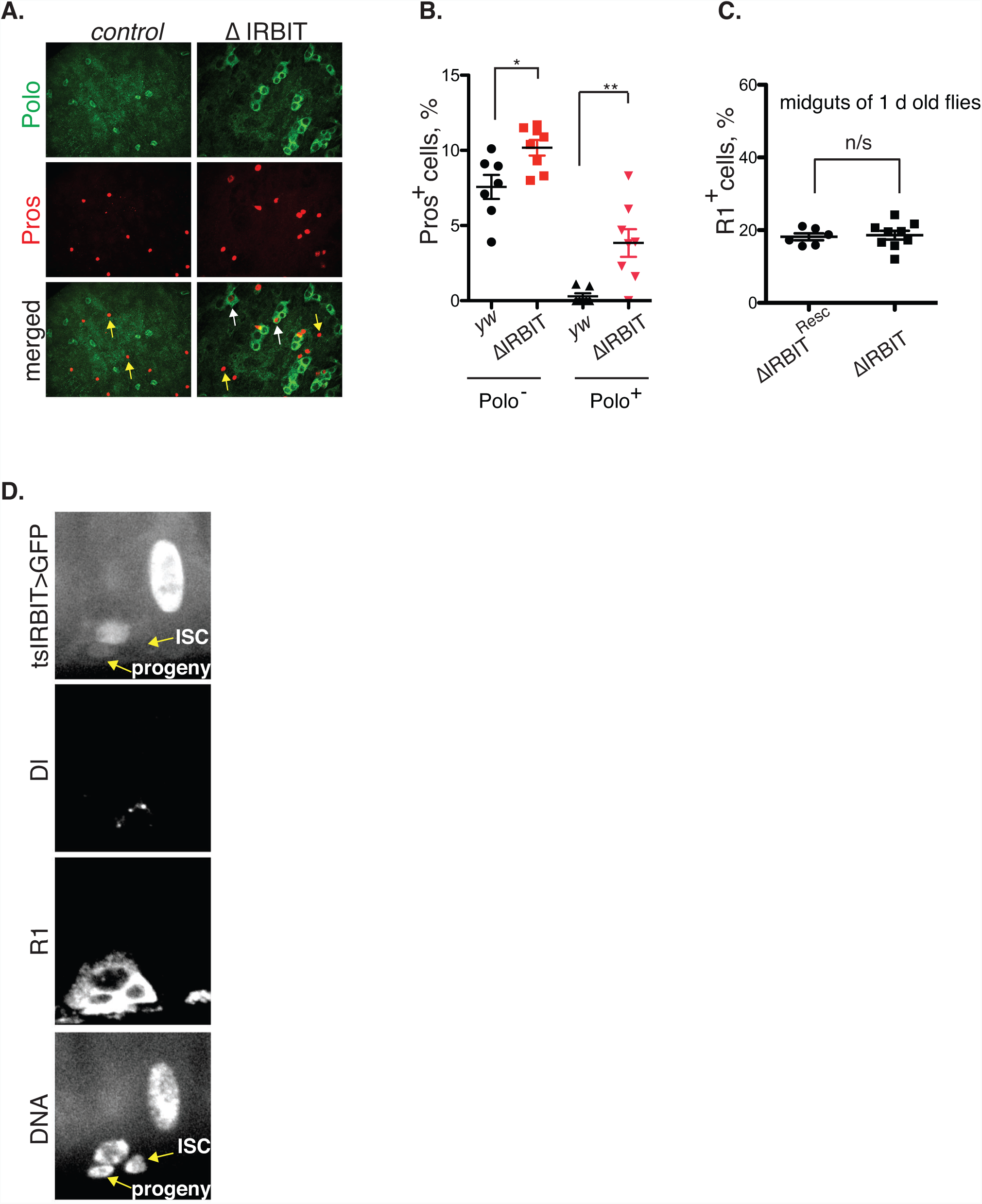
(A) ∆IRBIT mutant midguts have defects in the EEC cell differentiation. Midguts of 14 d old *yw*, ∆IRBIT and ∆IRBIT^Resc^ flies stained with antibodies against Pros (EEC lineage marker) and Polo. DNA was counterstained with Hoechst 33342. Note two types of Pros^+^ cells in the ∆IRBIT midguts: typical EEC Pros^+^ cells (yellow arrows) and Polo^+^/Pros^+^ cells (white arrows). Note, that only the regions of midguts with high numbers of EECs are shown. (B) Quantification of both types of EEC cells, as in (A). N=8 guts; error bars represent mean ± SEM. P values derived from unpaired t test with Welch’s correction, *P=0.017, **P=0.008. (C) Midguts of 1 d old (1d post eclosion) ∆IRBIT and ∆IRBIT^Resc^ flies were stained with R1 and DAPI and numbers of R1^+^ cells were calculated. N/s – not significant, two-tailed paired t-test. (D) Related to Figure 2E. Same image. A genomic construct that contains a putative *IRBIT* promoter and its 5’UTR was fused with *GAL80^ts^-P2A-GAL4* (^*ts*^IRBIT) and used to drive nlsGFP (*UAS-nlsGFP*) expression. Guts were dissected and co-stained with Dl (ISC marker) and R1. DNA was counterstained with Hoechst 33342. Note the presence of nuclear GFP in the ISC progeny (Dl-negative, R1-positive) but not in the ISC (Dl-positive, R1-positive). Both ISC and the progeny are indicated (arrows).

**Figure S5.**
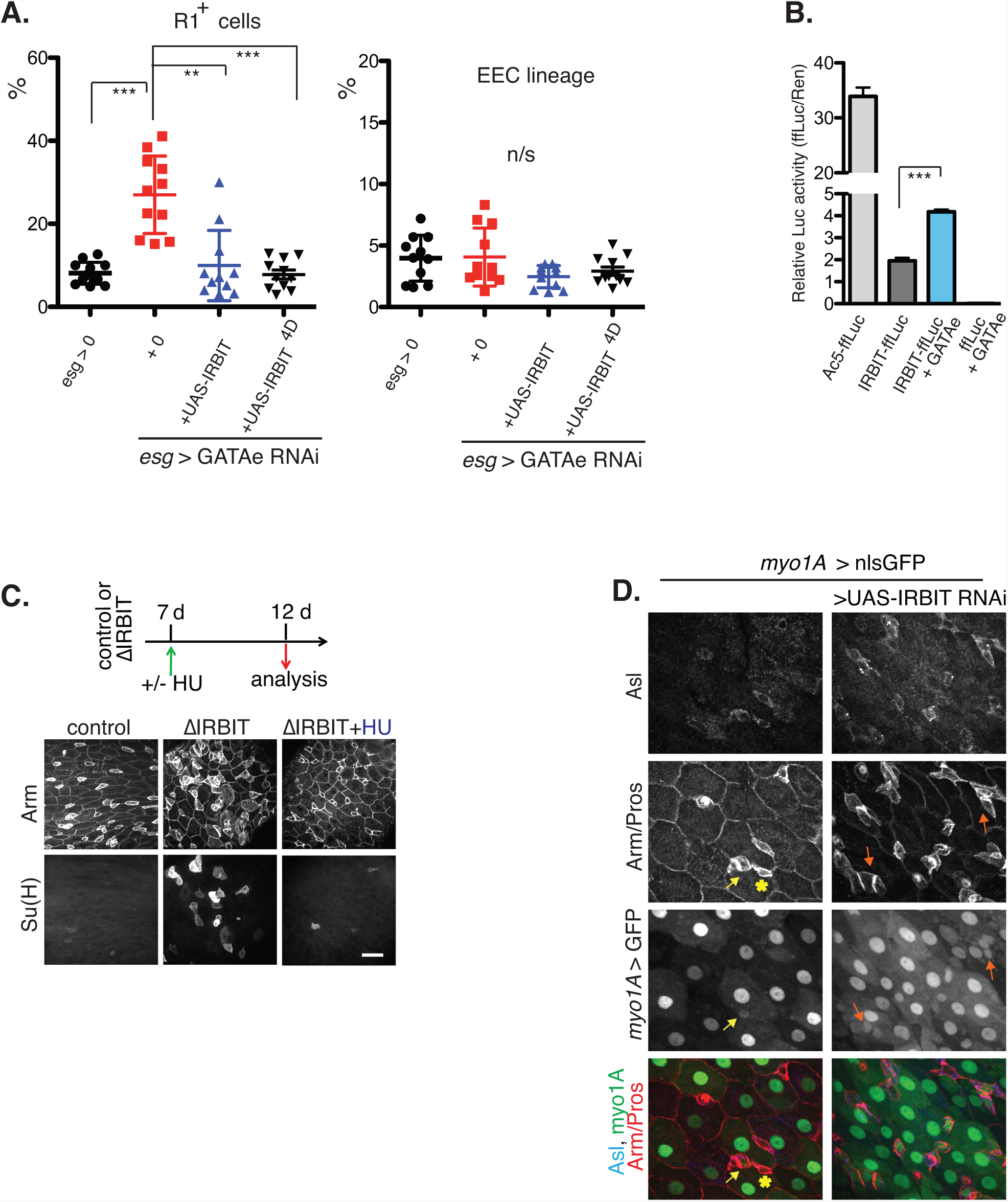
(A) IRBIT rescues GATAe deficiency. The expression of IRBIT was induced in GATAe-depleted ISCs and EBs (^*ts*^*esg, UAS-GATAe*^*RNAi*^, + *UAS-IRBIT or UAS-IRBIT*^*4D*^) for 5 d and R1^+^ or Pros^+^ cells were counted. Note that the depletion of GATAe is better rescued by phosphomimetic IRBIT mutant (IRBIT^4D^: ^55^SL<u>D</u>A<u>DDD</u>DSFSS) (Arnaoutov and Dasso, 2014) Kruskal-Wallis test with Dunn’s multiple comparison test. N=11; n/s – not significant. (B) GATAe stimulates IRBIT transcription. (C) Suppression of RNR activity rescues ∆IRBIT phenotype. 7 d old control and ∆IRBIT female flies, with their EB marked with ^*ts*^Su(H)>GFP were switched to food containing hydroxyurea (HU) for 5 days; midguts were then stained for Arm and DNA as above. Note the disappearance of Su(H)^+^ aggregates and the restoration of the normal gut architecture in the HU-treated ∆IRBIT guts. (D) Suppression of IRBIT transcription in EBs and ECs recapitulates ∆IRBIT phenotype. The expression of IRBIT was silenced for 5 d using *myo1A-gal4* driver. Note that myo1A is active in both ECs (large nuclei) and ISC progenitors (yellow arrow, Asl^+^ cell. Putative ISC is indicated with an asterisk). Note similarities between *myo1A*>*IRBIT^RNAi^* and ∆IRBIT (e.g. Figure 3F) and accumulation of Myo1A^+^, Asl^+^ cells (ISC progenitors) in ∆IRBIT.

**Figure S6.**
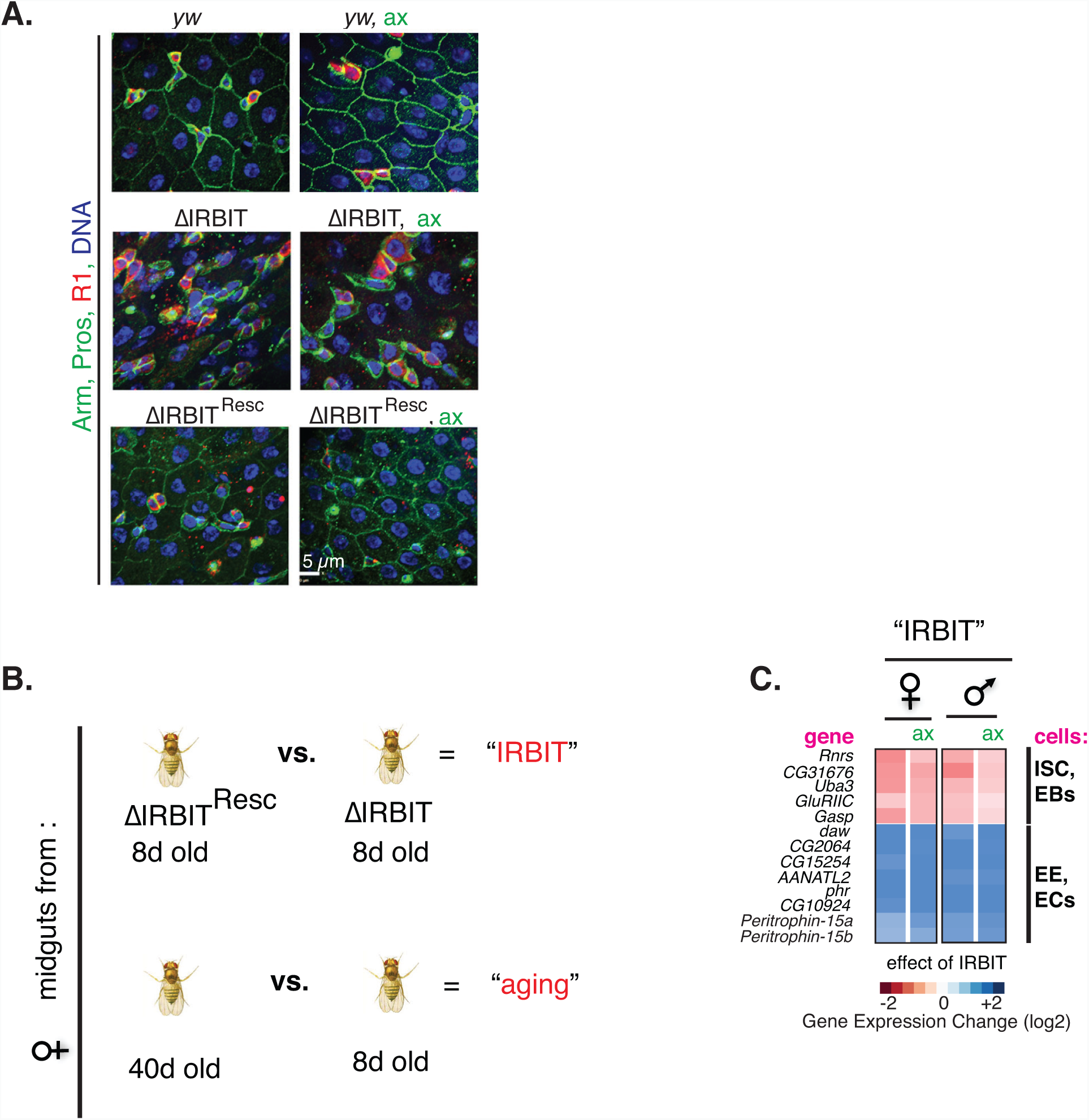
RNA-Seq profiling of midguts. (A) IRBIT-mediated differentiation of EB to EC is independent of microbiota. Midguts of 8 d old control (Hoskins et al.), ∆IRBIT and ∆IRBIT^Resc^ flies reared under normal or axenic (free of microorganisms, (ax)) conditions were stained with Arm and Pros (both in green), R1 (red) and DNA (blue). Note that ∆IRBIT flies have similar midgut morphology (R1^+^ cell clusters), independent of the intestinal load of microorganisms. (B) Schematics of the experiment for the RNAseq. (C) A list of DE IRBIT-dependent, microbiota- and sex-independent genes. Note that IRBIT causes both reduction of markers that belong to undifferentiated cells and increase of markers that belong to differentiated cells, indicating that IRBIT stimulates differentiation.

**Figure S7.**
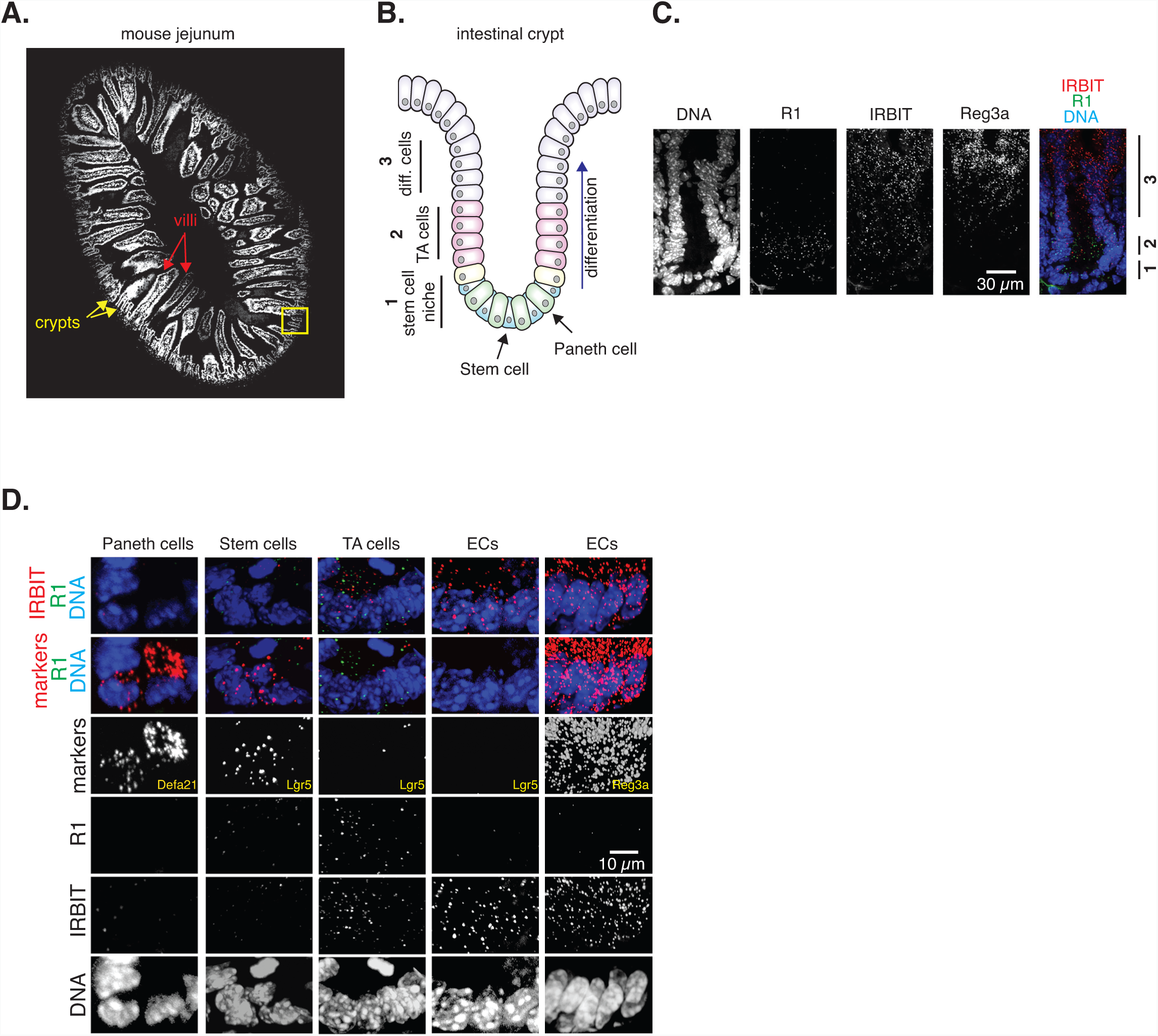
IRBIT mRNA expression pattern in mouse intestine. (A) Cross-section of mouse jejunum. (B) Scheme of the intestinal crypt. Positions of stem cells, Paneth cells, transiently amplifying (TA) cells and differentiating (diff.) cells are indicated. (C) Localization of R1, IRBIT and Reg3a transcripts. Note a gradient expression of IRBIT mRNA message, which coincides with the wave of differentiation. 1, 2, 3 – regions of the crypt, roughly corresponding to regions as depicted in (B). (D) Expression of IRBIT in specialized intestinal cells. ECs – differentiated enterocytes (in villi). Note that ECs (Lgr5-, Reg3a+) express IRBIT.

**Figure S8.**
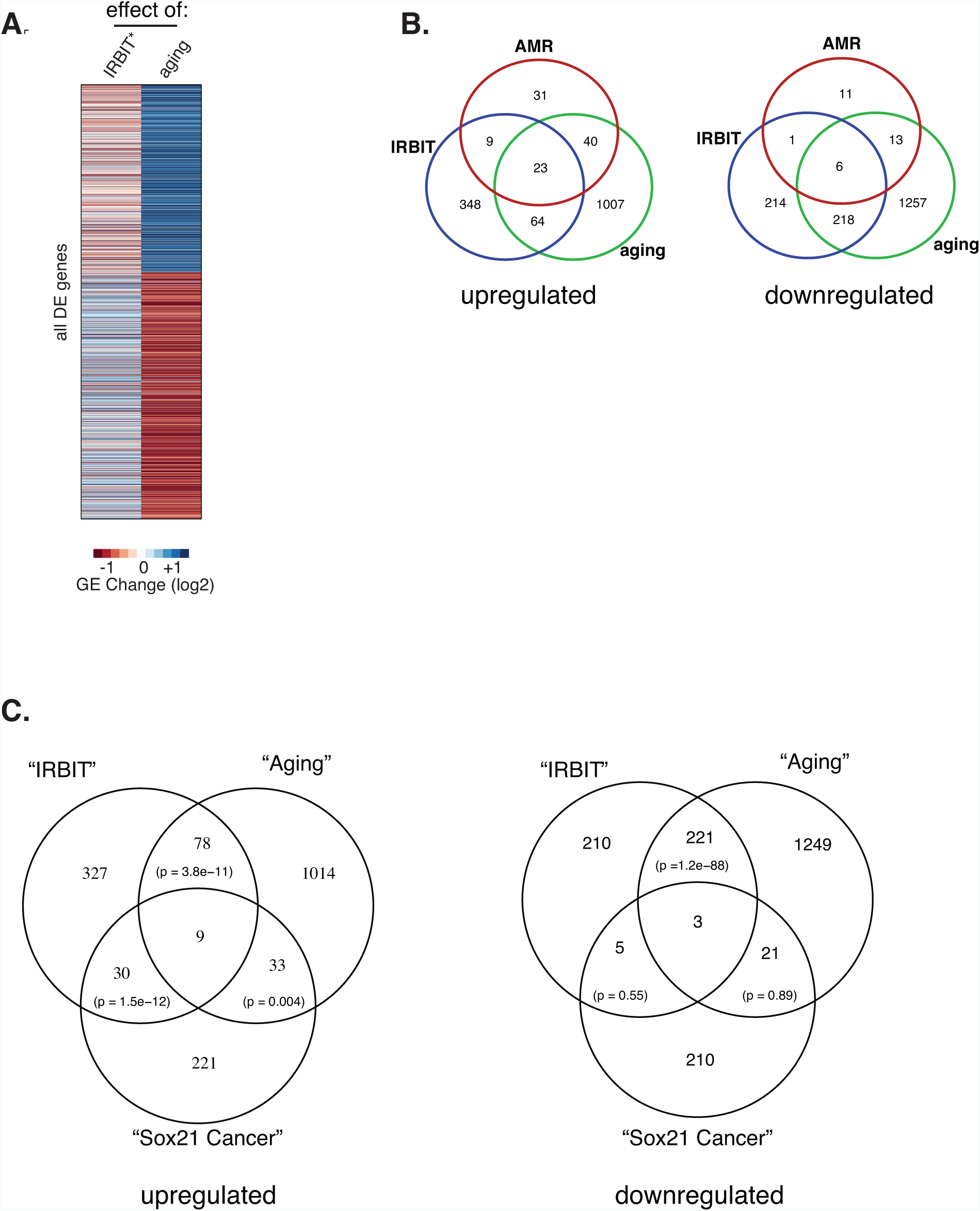
(A) The loss of IRBIT mirrors aging program in the gut. Clustering of “IRBIT*” (comparison of gene expression in 8d old midguts ΔIRBIT^Resc^ vs. ΔIRBIT) and “aging” (comparison of gene expression between *yw*, 40d old and *yw*, 8d old) DE genes. Note the anticorrelation of DE between “aging” and the effect of rescue of ∆IRBIT flies with genomic IRBIT construct (Pearson’s r = – 0.22, p < 2e-15, F test). (B) Increased anti-microbial response in ∆IRBIT midguts. Venn diagrams of the overlap between DE analysis, (as in Figure 4B). (C) ∆IRBIT midguts contain molecular signatures of stalled EBs. We obtained reprocessed count matrices (GSE117217), and performed differential expression analyses to compare DE of genes between EBs that bear Sox21a mutant and control EBs (we called the intersection of significant DE genes (adjusted p < 0.05) from the two studies (Chen et al., 2016; Zhai et al., 2017)) – “Sox21 cancer” – and used it to produce a Venn diagram to display the overlaps between the effects of Sox21a loss, aging and the loss of IRBIT (as in (A)). We used one-tailed hypergeometric test to describe the significance of overlap among gene sets.

**Figure S9.**
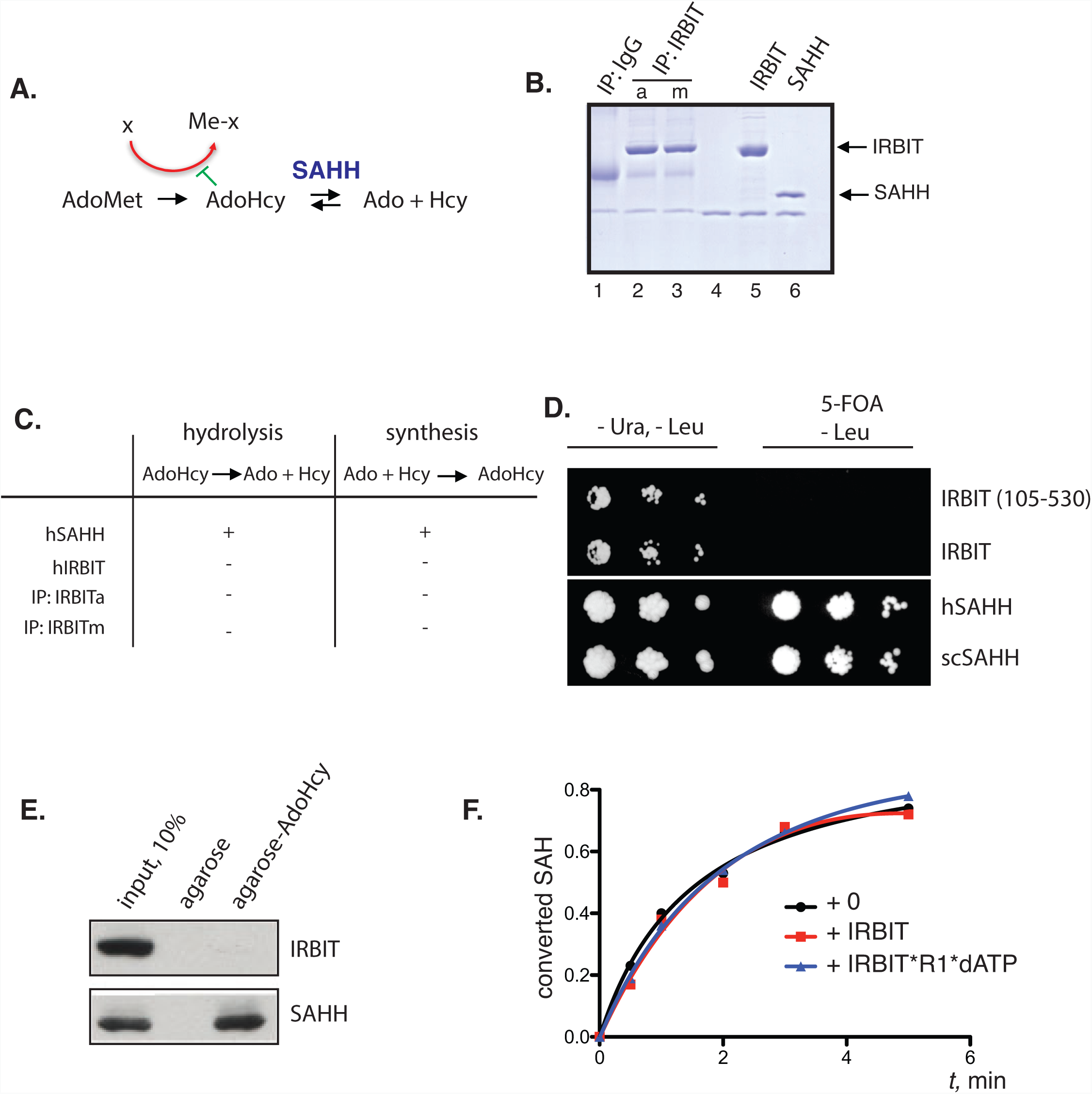
IRBIT is not a competitive inhibitor of SAHH. (A) SAHH-mediated reactions. AdoMet (S-adenosylmethionine), AdoHcy (S-adenosylhomocysteine), Ado (adenosine), Hcy (homocysteine). AdoMet is the source of methionine (Me) in methylation reactions. (B) IRBIT was immunoprecipitated from either asynchronous (a) or Nocodazole-arrested (m) HeLa cells (lanes 2, 3). IRBIT and SAHH were purified from baculovirus-infected SF9 cells (lanes 5, 6). Purified proteins were run on SDS-PAGE gel and stained with coumassie. (C) Proteins (as in (B)) were tested in indicated reactions. (D) IRBIT, SAHH-like domain of IRBIT, hSAHH and scSAH1 were tested for their capacity to rescue *sah1∆* yeast strain. (E) IRBIT does not bind AdoHcy. Recombinant hIRBIT and hSAHH (1 µg each) were purified on AdoHcy-agarose, and bound proteins were probed with antibodies against IRBIT and SAHH. (F) IRBIT is not a competitor of SAHH. Recombinant IRBIT or IRBIT*dATP*R1 complex were mixed with recombinant hSAHH (10:1 molar ratio) and assayed for AdoHcy hydrolysis.

**Table S1.** The list of differentially expressed IRBIT-, A+AMR-, and IRBIT^HU^ (HU)-dependent genes. DE is provided as fold changes, in log2 scale. Raw counts from the RNA-Seq analyses were also provided. Gene IDs and names are based on the FlyBase annotation.

**Table S2.** The list of differentially expressed IRBIT-, AMR-, and IRBIT^ax^ (axenic)-dependent genes.

**Table S2.** Related to Figure S8C. The list of differentially expressed IRBIT-, aging-, and Sox21a-dependent genes, with indicated overlaps.

